# Modulation of the mTOR pathway in male Lewis rats after morphine self-administration and subsequent extinction training

**DOI:** 10.1101/276121

**Authors:** Marcos Ucha, Santiago M Coria, Adrián E Núñez, Raquel Santos-Toscano, David Roura-Martínez, Javier Fernández Ruiz, Alejandro Higuera-Matas, Emilio Ambrosio

## Abstract

Addiction is a chronic disorder with an elevated risk of relapse, even after long periods of abstinence. Some of the neural mechanisms mediating addictions require protein synthesis, which could be relevant for the development of more effective treatments. The mTOR signaling pathway regulates protein synthesis processes that have recently been linked to the development of drug addiction. Thus, we have assessed the effects of morphine self-administration and its subsequent extinction on the expression of several genes that act in this pathway, and on the levels of some phosphoproteins in three brain areas related to reward learning and extinction: the amygdala, the nucleus accumbens, and the prefrontal cortex. We found an increase in *Raptor* and *Eif4ebp2* gene expression in the amygdala of rats that self-administered morphine, and this persisted beyond the extinction period. The expression of *Insr* in the amygdala of control animals decreased over time while the opposite effect was seen in the rats that self-administered morphine. We also found a strong correlation between some of the biochemical variables measured and behavioral traits, suggesting a significant role for the genes and phosphoproteins identified, mostly in the amygdala, in the behavioral effects of morphine.

## 1 Introduction

Addiction is a chronic debilitating condition with a high rate of relapse, for which as yet there is no effective treatment (Kalivas and O’Brien, 2008; McLellan et al., 2000). The mechanisms underlying the shift from controlled recreational use of drugs to pathological compulsive behavior are not yet fully understood, nor are the long-lasting neuroadaptive changes behind the elevated risk of relapse.

The development of an addiction depends on synaptic plasticity, which in turn relies on protein synthesis (Kalivas and O’Brien, 2008; Kauer and Malenka, 2007; Lüscher and Malenka, 2011). Thus, a signaling pathway that has generated much interest of late is that involving the mechanistic target of rapamycin, mTOR, a serine/threonine kinase that plays an important role in different aspects of cell growth, proliferation and survival (Kwon et al., 2003; Pearce et al., 2010; Zhou et al., 2009). This protein nucleates two different multi-protein complexes known as mTOR complex 1 (mTORC1) and mTOR complex 2 (mTORC2). These complexes are part of a pathway which integrates many intracellular and extracellular signals, and regulates processes such as protein, lipid and nucleotide synthesis (Düvel et al., 2010; Ma and Blenis, 2004; Porstmann et al., 2008; Stoica et al., 2011), autophagy (Blommaart et al., 1995), mitochondrial metabolism (Cunningham et al., 2007; Schieke et al., 2006) and cytoskeletal organization (Sarbassov et al., 2004). Given its role in protein synthesis-dependent synaptic plasticity (Casadio et al., 1999; Costa-Mattioli et al., 2009; Liu-Yesucevitz et al., 2011; Stoica et al., 2011), this pathway is thought to participate in the neurobiology of addictions. Accordingly, several studies have focused on the effects of rapamycin, an inhibitor of mTOR activity, on addictive behavior.

These studies suggest that this signaling pathway is involved in the long-lasting neuroadaptations that occur as addictive disorders progress (Dayas et al., 2012; Neasta et al., 2014). For example, rapamycin was able to reduce the rewarding effects of cocaine (Bailey et al., 2012; J Wu et al., 2011) and amphetamine (Narita et al., 2005) when measured in a conditioned place preference (CPP) test (Wang et al., 2010). Systemic rapamycin injections also reduced motivation for self-administered cocaine in rats as measured in a progressive ratio schedule of reinforcement (James et al., 2016). In addition, there was a reduction in cue induced reinstatement of cocaine seeking when rapamycin was injected directly in the core of the nucleus accumbens (NAcc) (Wang et al., 2010) and the *Mtor* gene expression was down-regulated in the ventral striatum of relapse-prone rats (Brown et al., 2011). Rapamycin also blocked nicotine-induced behavioral sensitization and activation by effectors of mTORC1. It has also been suggested that the dopamine receptor 1/mTOR complex 1-dependent plasticity is recruited following a first alcohol exposure and that it may be a critical cellular component of reinforcement learning (Beckley et al., 2016). In terms of opiates, chronic morphine decreases the soma size of dopaminergic cells in the ventral tegmental area (VTA), and neurotransmitter release by these cells, while increasing their excitability, events that are dependent on mTORC2 activity (Mazei-Robison et al., 2011). Activation of the mTOR pathway in the CA3 hippocampal region is necessary for the acquisition of morphine CPP in rats (Cui et al., 2010). Moreover, systemic inhibition of mTOR with rapamycin after re-exposure to a morphine paired compartment inhibits CPP in a dose dependent fashion, an effect that was replicated with cocaine and alcohol (Lin et al., 2014). Hence, mTOR may play a role in the reconsolidation of drug paired memories. Elsewhere, a single dose of rapamycin was able to reduce the craving elicited by drug related cues in human heroin addicts (Shi et al., 2009).

To date, we are unaware of any study that has used a self-administration protocol to study the effects of opioids on the mTOR signaling pathway in rodents. Here, we assessed the effects of morphine self-administration, followed by extinction training, on the mTOR pathway in male Lewis rats. For this purpose, we chose three brain areas known for their involvement in opioid reinforcement and extinction learning: the amygdala, the NAcc, and the prefrontal cortex (PFC). The expression of several mediators of the mTOR pathway was analyzed using RT-qPCR and we also assessed the levels of some of their phosphorylation sites required for their activation in western blots.

## 2 Methods

### 2.1 Animals

Adult male Lewis rats (Charles River Laboratories) were housed in groups of 4 in a temperature and humidity controlled facility, and on a 12h/12h light/dark cycle (lights on at 8:00am) with *ad libitum* access to food (standard commercial rodent diet A04/A03: Panlab) and water. Animals were allowed at least one week to acclimatize to the animal facility and they weighed around 250-300 g when the experimental procedures commenced. All the animals were maintained and handled according to European Union guidelines for the care of laboratory animals (EU Directive 2010/63/EU governing animal experimentation).

### 2.2 Apparatus

Twelve operant conditioning chambers (Coulborne Instruments), each equipped with a pellet dispenser and a microliter injection pump, were used to assess operant food rewarded behavior, and for the morphine self-administration and extinction studies. A catheter was connected to the rats and held in place with a spring-tether system, and a rotating swivel, which allowed the animals to move freely inside the chamber. Two levers were available throughout all the sessions, one of them inactive.

### 2.3 Experimental protocol

#### 2.3.1– Lever press instrumental training

At the beginning of the experiment, all the rats received daily instrumental training sessions with food pellets as reinforcers (grain-based rodent tablet, Testdiet™) on a fixed ratio 1 schedule, facilitating the acquisition of self-administration behavior. The sessions lasted 30 minutes and continued until the animals developed a robust lever press behavior (at least 100 lever presses in three consecutive training sessions).

#### 2.3.2. Surgery

Rats were anesthetized with an isoflurane/oxygen mixture, and a catheter was inserted into the right jugular vein of the animal and secured there with silk thread knots. The catheter was fixed subcutaneously around the neck, exiting the skin at the midscapular region. A pedestal of dental cement was then mounted on the skull of the rat in order to attach the tethering system. After surgery, the rats were allowed to recover for 7 days and a NSAID (meloxicam - Metacam™: 15 drops of a 1.5 g/ml solution per 500 ml of water) was added to the drinking water. Until the end of the self-administration procedure, the catheters were flushed daily with a sterile saline solution containing sodium heparin (100 IU/ml) and gentamicin (1mg/ml) to maintain catheter patency and to prevent infections.

#### 2.3.3. Morphine self-administration

A week after recovery from surgery, the rats underwent 19 daily sessions of morphine self-administration. During the dark phase of the light cycle, for 12 hours rats were allowed daily access to morphine (1 mg/kg in a sterile saline −0.9% NaCl-solution) or its vehicle alone under a fixed-ratio 1 reinforced schedule. During these sessions, one active lever press resulted in morphine infusion (1 mg/kg morphine in saline solution delivered over 10 seconds) followed by a 10 second time-out. A light cue located above the lever indicated the availability of the drug and a limit of 50 infusions per session was set in order to avoid overdosing. One day after the last session, two groups of rats were sacrificed (Vehicle self-administration - VhSA, n=10; Morphine self-administration - MSA, n=10), and their brains were processed and stored.

#### 2.3.4. Extinction training

The remaining rats were given 15 daily sessions of extinction training using the same self-administration protocol, although in this phase all the rats received a saline solution instead of morphine. One day after the last extinction session, the two remaining groups of rats (Vehicle extinction - VhEx, n=8; Morphine extinction - MEx, n=8) were sacrificed, and their brains processed and stored.

### 2.4 Sample processing

On the day of the sacrifice, the rats were decapitated and the amygdala (mainly the basolateral amygdala - BLA), NAcc (both shell and core) and PFC (mostly the orbitofrontal cortex, OFC) were dissected from the brains using a brain matrix (see Fig. 2). All the surfaces and tools used for dissection were sterilized and treated with RNAseZap^®^ (Ambion™), and all the steps were carried out with caution to maintain RNA integrity. The tissue samples from one hemisphere (randomized) were preserved overnight at 4 °C in RNAlater^®^ (Ambion™) and then stored at −70 °C in RNAlater^®^ for later RT-qPCR analysis. The samples of the other hemisphere were snap frozen with dry ice and stored at −70° for western blot analysis.

**Figure 1:**
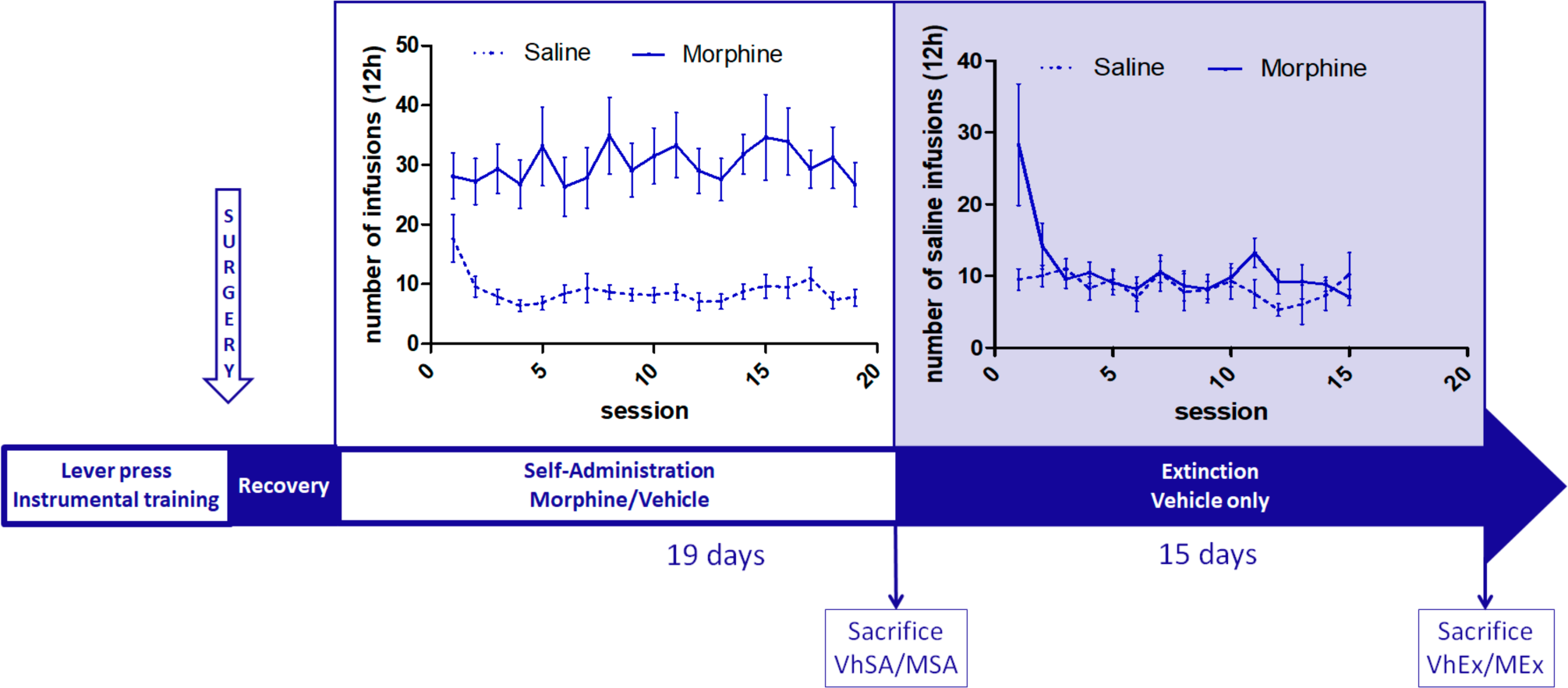
Timeline of the experimental procedures with a graphical representation of the behavioral data (Vehicle self-administration - VhSA, n=10; MSA - Morphine self-administration, n=10; VhEx - Vehicle extinction, n=8; MEx - Morphine extinction, n=8).

**Figure 2:**
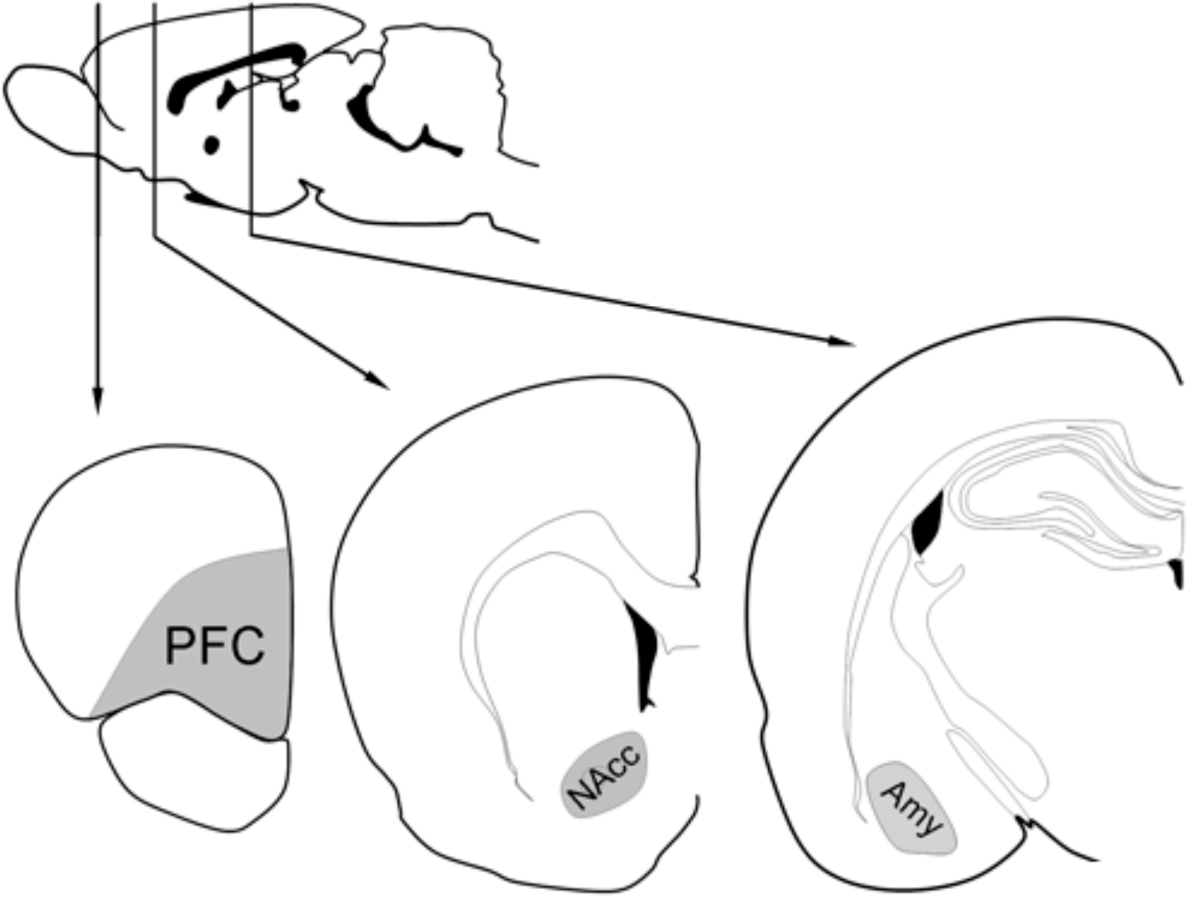
Schematic representation of the sections of the rat brain with the areas dissected out highlighted in gray.

### 2.5 RT-qPCR analysis

The samples stored in RNAlater^®^ were homogenized in QIAzol lysis reagent (QIAgen) using a pellet pestle. The total RNA was extracted and precipitated using the chloroform, isopropanol and ethanol method (Chomczynski and Sacchi, 1987) with glycogen as a carrier. The precipitate was dissolved in RNAse free water, and the concentration and RNA integrity was assessed in a bioanalyzer (Agilent 2100). The RNA concentration in each sample was adjusted by adding RNAse free water and to avoid genomic DNA contamination, DNAse digestion was performed (DNAse I, Amplification Grade, Invitrogen) following the manufacturer’s instructions. Finally, the samples were retrotranscribed using a commercial kit (Biorad iScript™ cDNA Synthesis Kit). PCR assays were performed on a real time PCR detection system (CFX9600, Biorad) with a SSO Advanced SYBR mix (Biorad) using the primers indicated in the supplementary materials section. The raw data was normalized using the ΔΔCt method (Livak and Schmittgen, 2001) using *Gapdh* as a reference gene and the reaction efficiencies were obtained using LinRegPCR software (Ruijter et al., 2009).

### 2.6 Western blotting

The tissue samples were homogenized using a pellet pestle in 10 volumes of lysis buffer: 50mM HEPES [pH7.5], 320 mM sucrose, (Complete^™^ EDTA-free, Roche) protease inhibitors, and phosphatase inhibitors (PHOStop^™^, Roche). The resulting homogenate was centrifuged at 2000 g and at 4 °C for 10 minutes, the supernatants were recovered and their protein concentration was assessed using the Bradford assay (Bio-Rad Protein Assay). The protein extracts (3 mg) were mixed with 6X Laemmli buffer and loaded onto 8% SDS-PAGE gels, resolved by electrophoresis and transferred to PVDF membranes. After blocking non-specific interactions with 5% BSA for one hour, the membranes were probed overnight with the primary antibodies (see supplementary materials) that were then recognized with a horseradish peroxidase-conjugated secondary antibody (see supplementary materials). Antibody binding was visualized by chemiluminescence (ECL Plus Western Blotting Substrate, Pierce™). As a control for protein loading, we measured the total protein loaded by adding 2,2,2-trichloroethanol to the gels prior to polymerization (final concentration 0.5% v/v: Ladner et al., 2004), and after resolving the gel, it was excited with an UV transilluminator and the fluorescence emitted was measured. We used a CCD based detector (Amersham Imager 600) to capture both the chemiluminiscence and the UV/fluorescence images, and the ImageJ software to analyze and quantify them. When necessary, antibodies were stripped using a harsh stripping protocol (“Stripping for reprobing”: Abcam^®^).

### 2.7 Statistical analysis

The data obtained from the behavioral experiments were analyzed using repeated measures ANOVA. The analysis of the self-administration data had Sessions as a within-subject factor, and Treatment (M or Vh) and Phase (SA or Ex) as the between-subject factors. To analyze the extinction behavioral data, only Treatment was used as the between-subject factor. The degrees of freedom were adjusted when the sphericity assumption was violated by applying the Greenhouse-Geisser correction.

To analyze the biochemical assays we performed two-way ANOVAs with two between-subject factors: Treatment and Phase. When the required assumptions for ANOVA were not met, we used a Kruskal-Wallis test followed by a multiple comparison of groups, based on the descriptions given by Conover (Conover, 1999).

Finally, we analyzed the potential correlations between the behavioral and biochemical variables, for which we selected the following dependent behavioral variables: cumulative consumption in the morphine self-administration sessions (TotSA), cumulative consumption in the last 5 morphine self-administration sessions (Last5SA), cumulative consumption during the first 3 extinction sessions (First3Ex); and the linear regression slope of the consumption during the first 3 extinction sessions (Slope3EX). We chose the first 3 extinction sessions because the number of lever presses of the group MEx from the third session was already similar to the VhEx group. We then measured the relationship between these variables and the biochemical variables using Pearson’s r, Spearman’s ρ and Kendall’s τ as the correlation coefficients. Given the heterogeneity of the distributions and the number of correlations studied, we only report those that were significant when assessed by the three methods indicated.

### 2.8 Software

All the statistical analyses were performed using SPSS 24 (IBM) and the level of significance was set to α=0.05. The Kruwal 10.3 software was used for the non-parametric multiple comparison of groups. All the graphs were designed using the PRISM 6 software (GraphPad Software, Inc).

## 3 Results

### 3.1 Behavioral data

All the experimental animals achieved a high number of lever presses during the acquisition phase, probably due to the previous autoshaping training (Fig. 1). Subsequently, the rats that received saline lowered the rate of lever pressing, whereas the number of lever presses of the rats that received morphine remained high. During the first extinction session, there was a surge in the number of lever presses in the rats of the MEx group, although this decreased gradually in the following sessions until it reached values similar to those of the VhEx group. When we performed the Mauchly’s test for sphericity, the self-administration data did not meet the assumption of sphericity (χ^2^_170_=343.11, p<0.001) and therefore, the degrees of freedom were corrected using a Greenhouse-Geisser transformation (ε=0.45). The two way-repeated measures mixed ANOVA showed a significant effect of the Sessions factor (F_8.03,248.89_=4.23, p<0.001, ηG^2^=0.05) and an almost significant Treatment x Sessions interaction (F_8.029,248.891_=1.948, p=0.053, ηG^2^=0.024). Given that the Greenhouse-Geisser transformation employed is particularly conservative and considering the lever presses profile observed (see Fig. 1), the lever presses during the sessions are likely to be different for the MSA and VhSA groups. Indeed, when we assessed the overall average values of all the self-administration sessions we found a significant effect of the Treatment factor (F_1,31_=45,07, p<0.001, *η*^2^=0.98). We did not find any significant Treatment*Phase interaction (F_1,31_=10,43, p>0.05) or any effect of the Phase factor (F_1,31_=84,63, p>0.05). Therefore, it was concluded that the groups that underwent extinction performed similarly to their counterparts during the self-administration procedure. Regarding the extinction session data, the assumption of sphericity was also violated (χ^2^_104_=198.23, p<0.001) and a Greenhouse-Geisser correction had to be applied again (ε=0.26). Similarly, we found a significant effect of the Sessions factor 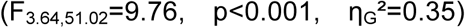 and a significant Treatment x Session interaction 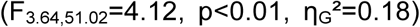, showing that both experimental groups performed differently across the sessions. We also found a significant effect of the Treatment factor (F_1,14_=10.99, p<0.01, *η*^2^=0.92) for the average values throughout the extinction sessions (see Fig. 1). To test whether the rats in the MEx group had extinguished the morphine self-administration behavior, we compared the mean number of lever presses during the last three days of extinction in the MEx and VhEx groups. Importantly, no significant differences were observed between these groups of rats (t_14_=-1.41, p>0.05).

### 3.2 Gene expression

In the amygdala, the gene expression analysis identified a significant effect of the treatment on the expression of the Regulatory Associated Protein of MTOR Complex 1 (*Rptor*) (F_1,28_=5.57, p<0.05, *η*^2^=0.16) and the Eukaryotic Translation Initiation Factor 4E Binding Protein 2 (*Eif4ebp2*) (F_1,28_=4.28, p<0.05, *η*^2^=0.13: Table 1). The expression of these genes increased in the rats that self-administered morphine and this effect persisted even after extinction training. In this structure, we also found a main effect of the Phase factor on the expression of AKT Serine/Threonine Kinase 1 (*Akt1*) (F_1,28_=6.9, p<0.05, *η*^2^=0.19) and the Insulin Like Growth Factor 2 Receptor (*Igf2r*) (F_1,28_=5.74, p<0.05, *η*^2^=0.15). In both cases transcription was enhanced after the extinction sessions. Significant differences in the Insulin Receptor (*Insr*) expression were evident between the four experimental groups 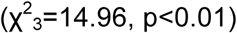 and the multiple comparison test showed that the VhSA rats expressed *Insr* more strongly than the MSA and VhEx rats. *Igf2r* expression was also affected In the PFC by the Phase factor (F_1,26_=7.32, p<0.05, *η*^2^=0.21), although its expression was weaker after the extinction sessions. The Treatment x Phase interaction appeared to affect the expression of *Eif4ebp2* (F_1,26_=4.18, p=0.14, *η*^2^=0.04) but after the simple effects analysis, this effect appeared not to be significant. There were no statistically significant differences in the expression of any of the genes analyzed in the NAcc.

**Table 1:**
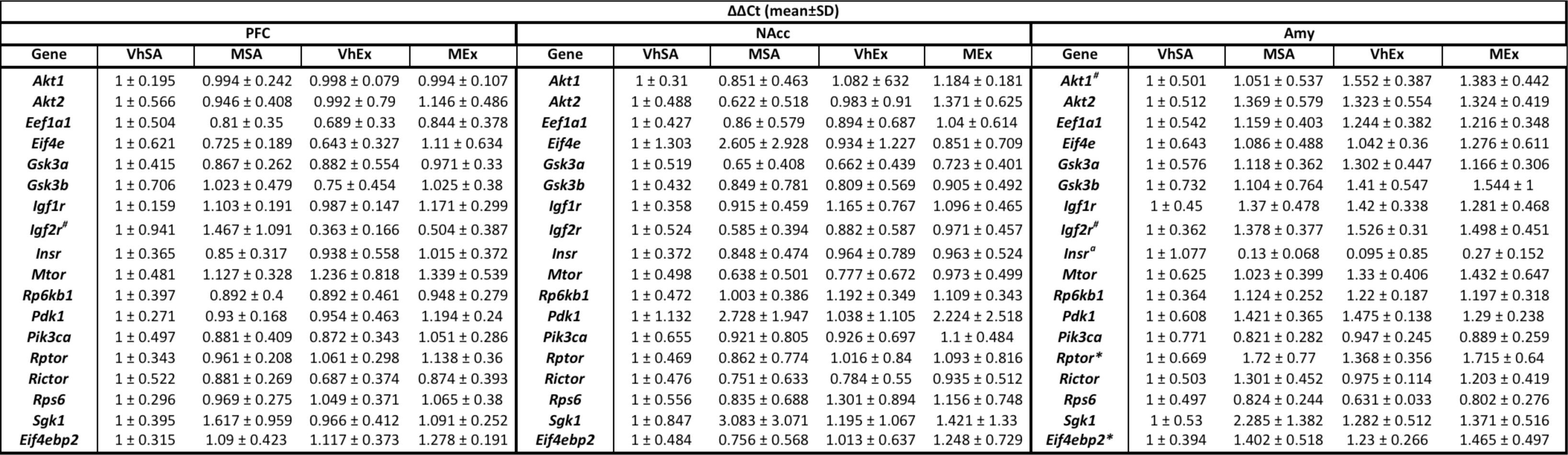
Results of the gene expression data expressed as the mean ± SD of the ΔΔCt value: * Effect of the treatment; ^#^ Effect of the phase; ^a^ VhSA > VhEx&MSA.

### 3.3 Phosphoprotein levels

We did not find any significant effects of the Treatment on the phosphoproteins assessed in each of the brain areas examined. However, in the amygdala the Phase factor affected the levels of phospho-GSK-3α (Ser21/9) (F_1,28_=5.32, p<0.05, *η*^2^=0.14) and the 68kDa band of phospho-PDK1 (Ser241) (F_1,29_=6.18, p<0.05, *η*^2^=0.17). The levels of both these phosphoproteins were lower after the extinction sessions (Fig. 3,4 and 5).

**Figure 3:**
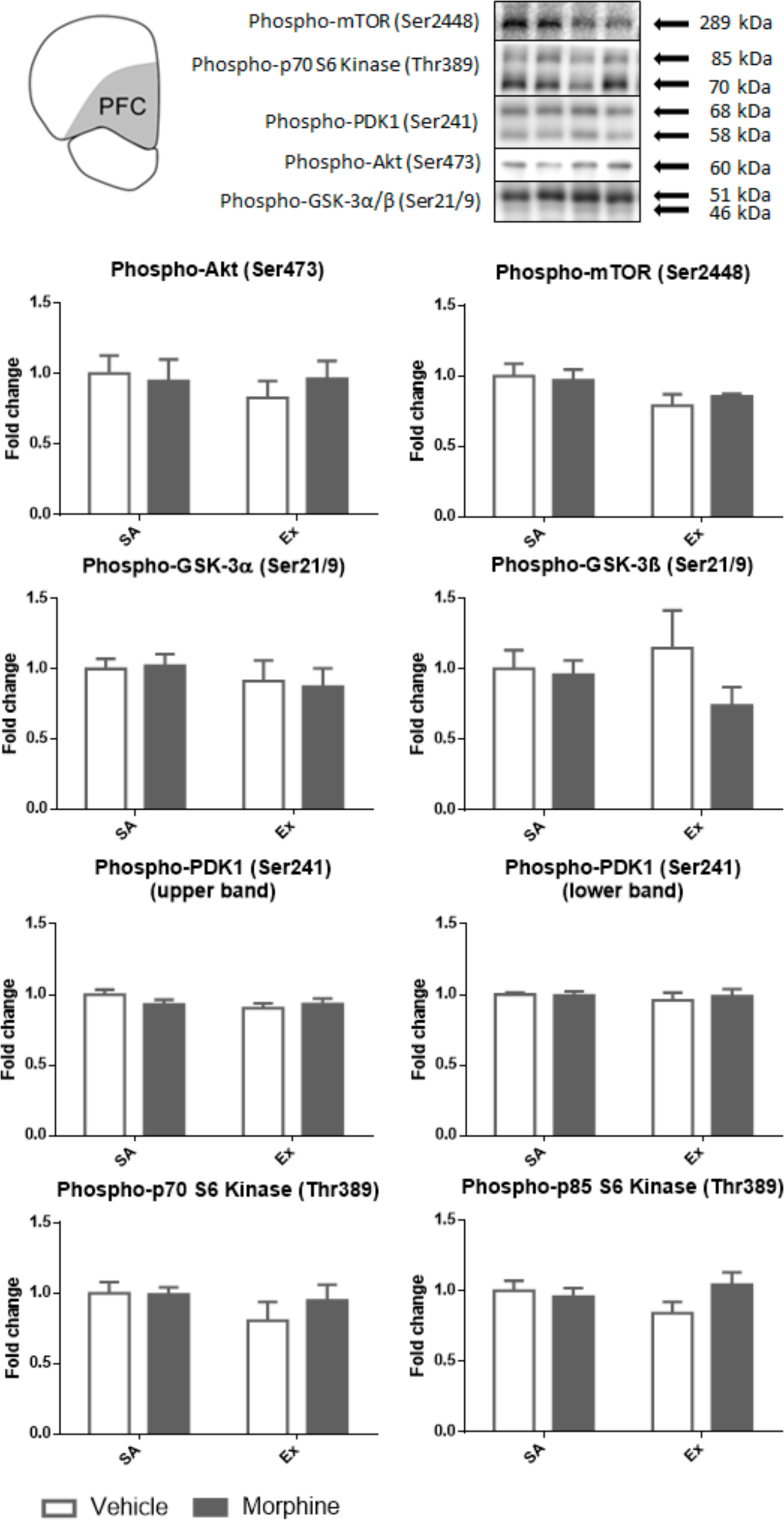
Representative Western Blots to analyze phosphoproteins in the PFC, normalizing the data to the total protein in the gel and the mean of the VhSA group (expressed as the mean ± SD).

**Figure 4:**
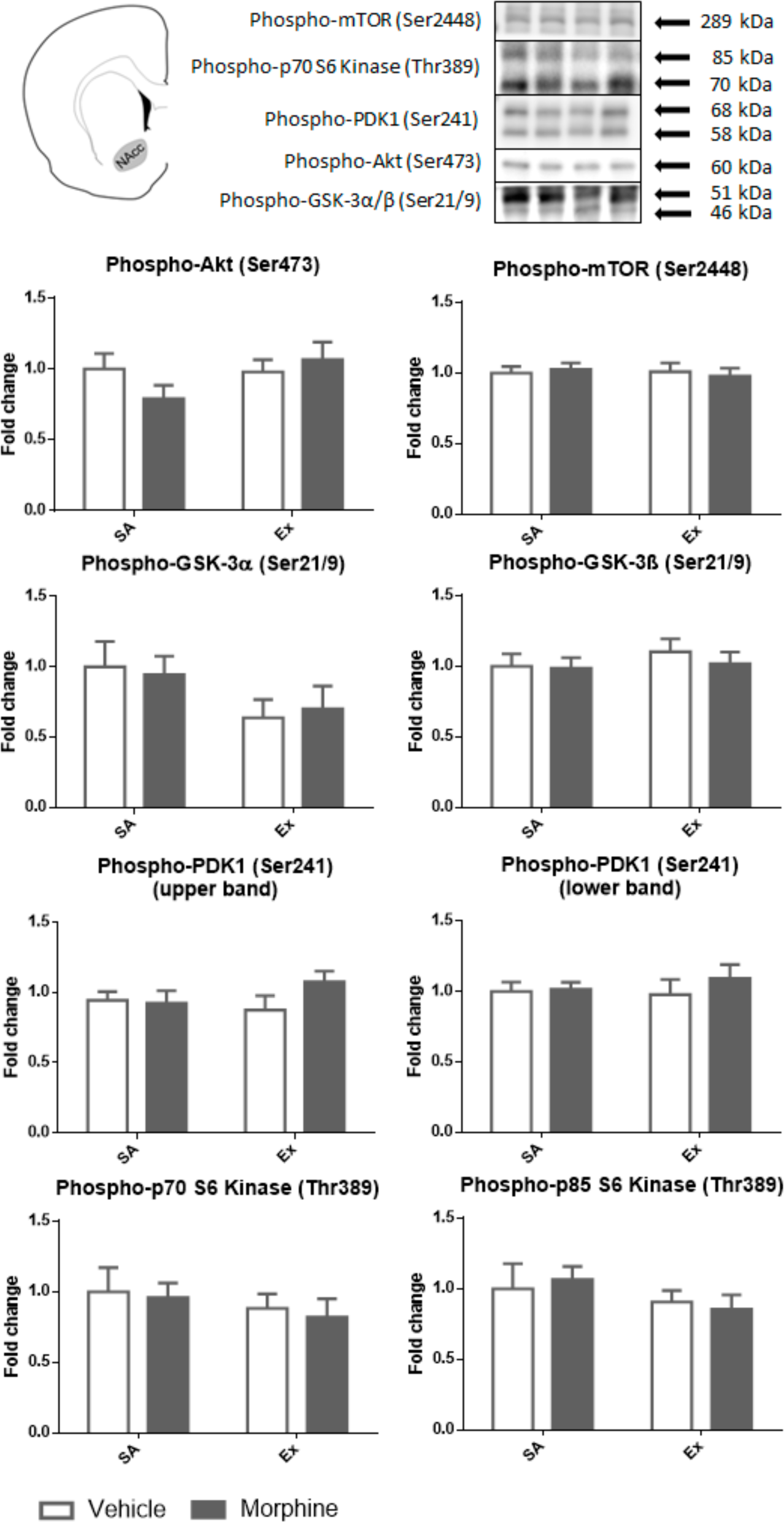
Representative Western Blots to analyze phosphoproteins in the NAcc, normalizing the data to the total protein in the gel and the mean of the VhSA group (expressed as the mean ± SD).

**Figure 5:**
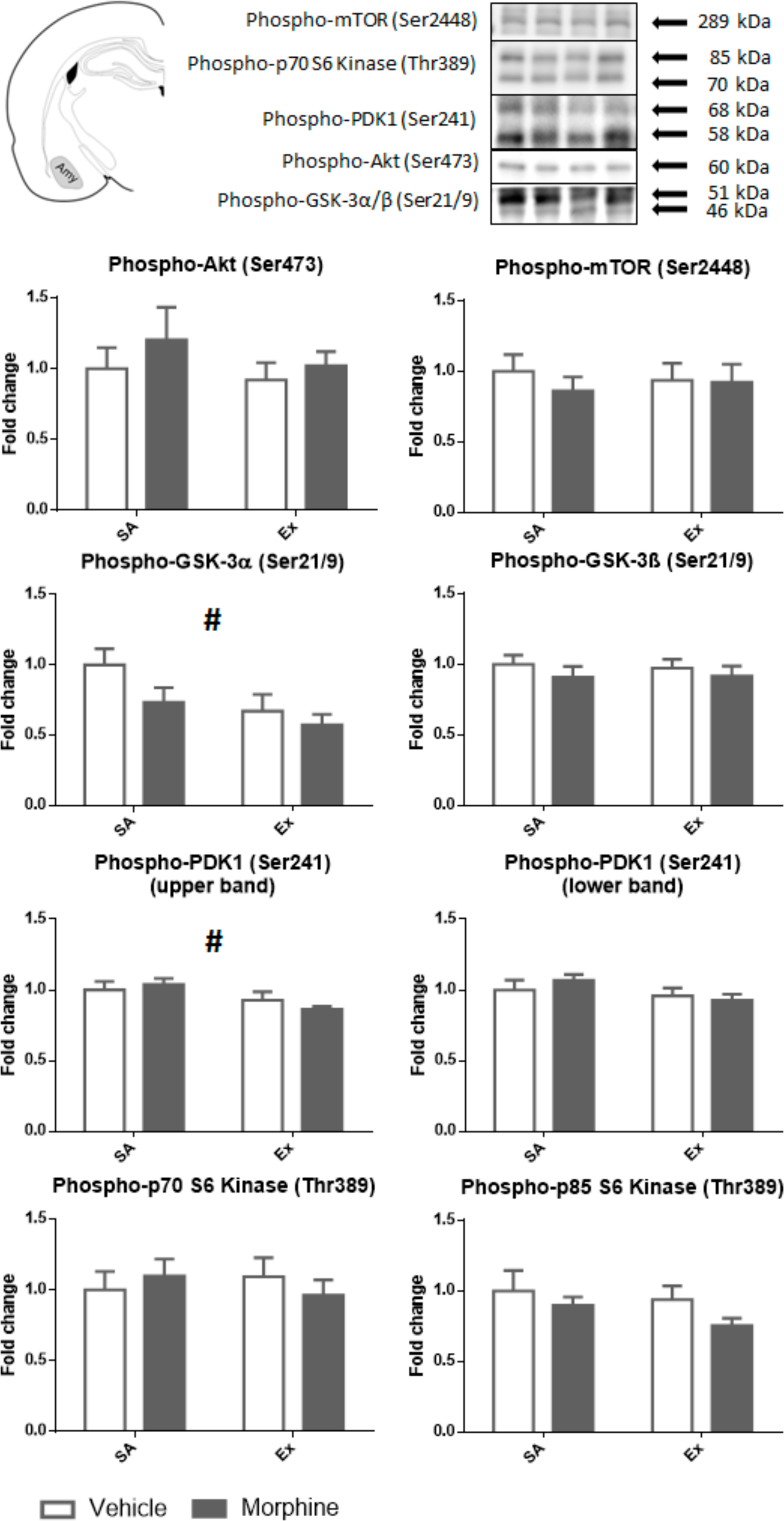
Representative Western Blots to analyze phosphoproteins in the Amy, normalizing the data to the total protein in the gel and the mean of the VhSA group (expressed as the mean ± SD). # Effect of the phase.

### 3.4 Correlations

In the MSA group (Fig. 6), the TotSA was correlated with the expression of the Eukaryotic Translation Elongation Factor 1 Alpha 1 (*Eef1a1*) in the amygdala (r=0.86, p<0.01; ρ=0.79, p<0.05; T=0.571; p<0.05). Likewise, in these rats the Last5SA was correlated with the expression of *Eef1a1* (r=0.77, p<0.05; ρ=0.81, p<0.05; T=0,64, p<0.05) and the Serum/Glucocorticoid Regulated Kinase 1 (*Sgk1*) (r=0.76, p<0.05; ρ=0.71, p<0.05; τ=0.571, p<0.05) in the amygdala. In the MEx group (Fig. 7), we found that *Mtor* expression in the amygdala was significantly correlated to Last5SA (r=0.82, p<0.05; ρ=0.88, p<0.01; p=0.79, p<0.01) and that Glycogen Synthase Kinase 3 Alpha (*Gsk3a*) mRNA expression in the amygdala was correlated to the First3Ex (r=0.79, p<0.05; ρ=0.71, p<0.05; τ=0.76, p<0.01). Similarly, there was also a significant inverse correlation between both the upper (r=-0.92, p<0.01; ρ=-0.75, p<0.05; τ=-0.62, p<0.05) and lower (r=0.81, p<0.05; ρ=0.8, p<0.05; τ=-0.69, p<0.05) phPDK1 (Ser241) bands in the PFC and Slope3Ex.

**Figure 6:**
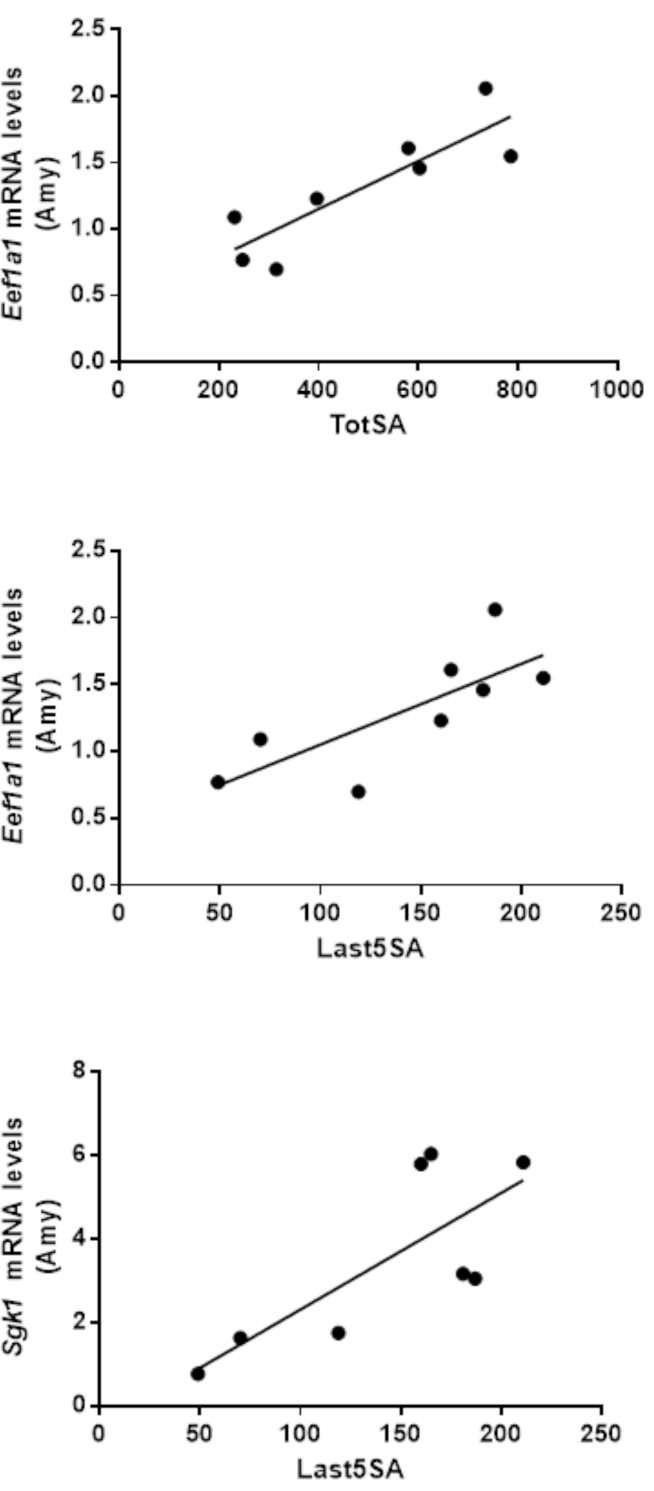
Significant correlations between the biochemical and behavioral variables in the MSA group.

**Figure 7:**
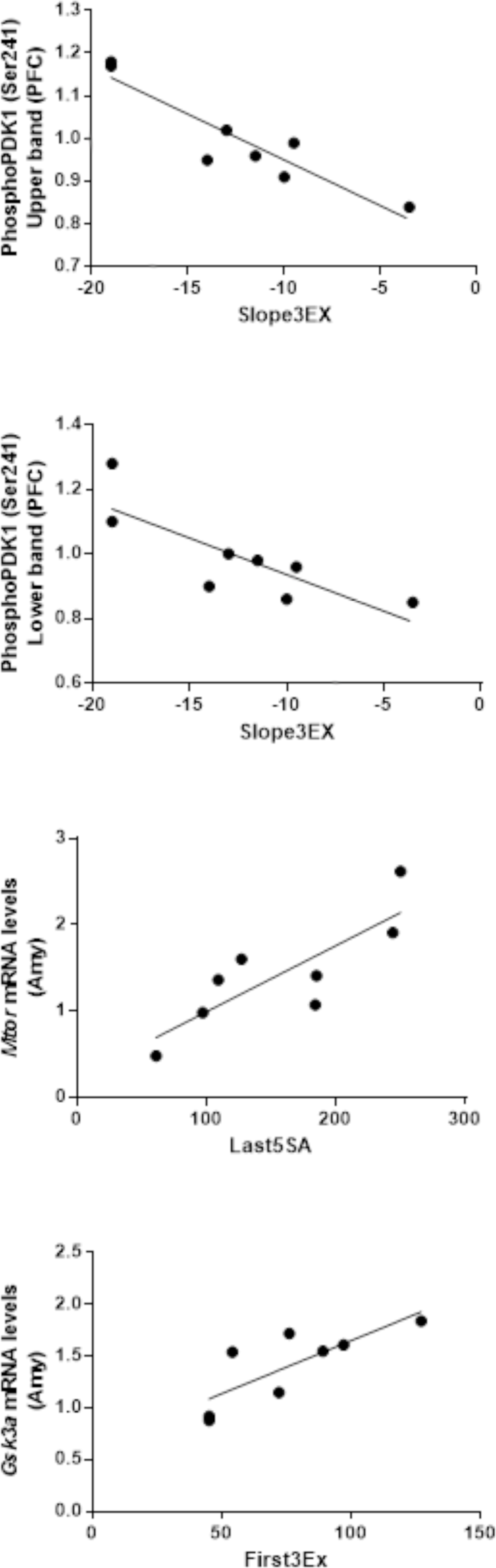
Significant correlations between the biochemical and behavioral variables in the MEx group.

## 4 Discussion

We assessed the effects of morphine self-administration and the subsequent extinction of this behavior on the expression of several genes and on the levels of some phosphorylated proteins of the mTOR signaling pathway in three brain areas related to reward learning and extinction: the amygdala, the NAcc and the prefrontal cortex. The morphine self-administration program employed only affected the expression of the *Rptor* and *Eif4ebp2* genes in the amygdala, an effect that persisted after extinction (Table 1). The *Rptor* gene encodes the regulatory-associated protein of mTOR (Raptor), a protein in the mTOR complex 1 (mTORC1), while the product of the *Eif4ebp2* gene is the eukaryotic translation initiation factor 4E-binding protein 2 (EIF4EBP2), one of the downstream effectors of this complex (Shimobayashi and Hall, 2014). Raptor regulates mTOR kinase activity, and it also recruits mTORC1 substrates like the S6 kinases and EIF4E binding proteins like EIF4EBP2 (Hara et al., 2002; Kim and Sabatini, 2004; Ma and Blenis, 2009). The eIF4EBP proteins in turn regulate EIF4E activity, which is responsible for the cap-dependent translation of mRNAs (Richter and Sonenberg, 2005). Our dissection of the amygdala mostly included the BLA, an area with an important role in conditioning learning given that it encodes the motivational value of the conditioned stimulus, either appetitive or aversive (Everitt et al., 2003). The BLA also has a role in the formation, retrieval and reconsolidation of drug-related memories (Luo et al., 2013). Indeed, c-Fos activity in the BLA is enhanced in rats showing CPP or conditioned place aversion (CPA) to morphine (Guo et al., 2008). Considering all this evidence together, the enduring increase in mTORC1 activity after morphine self-administration in the BLA (as suggested by the elevated transcription of the *Rptor* and *Eif4ebp2* genes) could contribute to the stabilization of those morphine-related aversive and appetitive memories that persist even after extinction.

Another interesting result was the variation in *Insr* gene expression that decreases drastically after morphine self-administration relative to rats exposed to the vehicle alone (Table 1). The *Insr* gene encodes the insulin receptor, one of the upstream activators of the PI3K/Akt/mTOR pathway (Niswender et al., 2003; Taha and Klip, 1999). Moreover, morphine can also activate this pathway through μ opioid receptors (Law et al., 2000; Polakiewicz et al., 1998). It is plausible that our results could reflect the opioid inhibition of insulin signaling due to a crosstalk between the downstream signaling pathways of both receptors, as shown previously in cell cultures (Li et al., 2003). These results are also consistent with the evidence that a chronic morphine regime downregulates the insulin receptor substrate 2 (IRS2)-Akt signaling pathway in the ventral tegmental area (Russo et al., 2007). This dampened endogenous insulin signaling might contribute to the development or expression of morphine withdrawal syndrome. Indeed, insulin administration reduces withdrawal symptoms in rats (Singh et al., 2015). Furthermore, rats that self-administered morphine did not display the decrease over time that vehicle treated rats did. This increase in the *Insr* might suggest recovery from withdrawal syndrome although direct evidence for this is lacking.

We also found some changes independent of the treatment but that rather reflected the experimental phase. The *Akt1* and *Igfr2* genes were more strongly expressed in the amygdala in the groups that underwent extinction training, even in the rats that received a saline solution during the self-administration phase. As opposed to the amygdala, *Igfr2* expression in the PFC was reduced in both groups after extinction (Table 1). These changes could reflect the natural regulation of these genes over the lifetime of the rats or maybe, they were a result of the experimental manipulations the rats were subjected to (surgery, handling, behavioral experiments…). Apart from the changes in gene expression, we also found variations in the phosphorylation of GSK-3α (Ser21/9) and of the 68kDa isoform of PDK1 (Ser241), both of which changed after extinction in the two groups irrespective of their prior treatment (Fig. 5). The levels of both phosphoproteins decreased in the BLA after extinction, and those of phospho-GSK-3α (Ser21/9) also tended to fall in the NAcc (Fig. 4).

Although most of the variables measured did not differ significantly between the experimental groups, some were correlated with behavioral variables and this leads to some interesting interpretations. For example, the *Eef1a1* mRNA transcripts in the amygdala of the MSA rats were positively correlated with overall morphine consumption, and with its consumption during the last 5 days of self-administration (Fig. 4). The *Eef1a1* gene encodes the Elongation factor 1-alpha 1 protein (eEF1a1). This protein is responsible for the enzymatic delivery of all aminoacyl-tRNAs to the ribosome, permitting the elongation of growing peptides and making it a key element integrating the cellular pathways regulating protein synthesis (Mateyak and Kinzy, 2010). *Eef1a* translation is supposed to be regulated by S6 phosphorylation by p70S6k, a mTORC1 target (Jefferies et al., 1994; Kimball et al., 1999). These findings, together with the long-lasting increased amygdalar expression of the *Rptor* and *eif4ebp2* genes in morphine-exposed animals, suggest that this opiate activates the translational machinery in the amygdala. As suggested previously, such an effect could be indicative of the engagement of plasticity-related process in a similar way to those invoked by long-term potentiation (Tsokas et al., 2005). We also found a positive correlation between the amygdalar *Sgk1* mRNA levels (serum and glucocorticoid-induced protein kinase 1) and morphine consumption during the last 5 days of self-administration (Fig. 4). Moreover, SGK1 mRNA expression was increased after morphine self-administration in all of the brain areas studied, although these changes didn´t reach statistical significance (Table 1). SGK1 is a serine/threonine kinase that is activated by the phosphorylation of a hydrophobic motif at Ser^422^, a phenomenon mediated by the mTORC2 complex (García-Martínez and Alessi, 2008). Although we do not detect changes in proteins associated to the mTORC2 complex, such as Rictor, this correlation suggests the activity of this complex may be modulated by morphine consumption. To our knowledge, this is the first study showing the potential modulation of *Sgk1* by opiates in the amygdala, although other groups studying other brain areas have documented a similar effect. For example, *Sgk1* mRNA expression was enhanced in whole brain lysates after chronic oxycodone administration, a μ opioid receptor agonist (Hassan et al., 2009). Elsewhere, *Sgk1* mRNA levels and activity was seen to increase in the VTA after 7 days of passive morphine administration (i.p. 15mg/kg: Heller et al., 2015) and chronic morphine administration passively increases mTORC1 activity in the VTA, while decreasing that of mTORC2. Such treatment also decreased the soma size of VTA dopaminergic neurons, an effect that increased cell activity but that decreased dopamine output in the NAcc shell. These effects were blocked by overexpressing *Rictor* in the VTA, indicating that reduced mTORC2 activity mediates these adaptations (Mazei-Robison et al., 2011). SGK1 has previously been shown to play an important role in spatial memory consolidation (Lee et al., 2006; Tsai et al., 2002) and LTP (Ma, 2006), and in the light of these results, this kinase may also be important in the effects of opioids in the brain as well.

We also found a correlation between *Mtor* expression in the amygdala and the consumption of morphine during the last five days of self-administration, yet only in the MEx rats (Fig. 5). The BLA is known to participate in the memory consolidation processes underlying extinction, as evident in cocaine extinction (Fuchs et al., 2002), extinction of fear (Falls et al., 1992; Lu et al., 2001) and extinction of amphetamine CPP (Schroeder and Packard, 2004). The fact that we found this correlation only after extinction training could mean that *Mtor* expression in the amygdala has a role in the consolidation of extinction memory. Indeed, the inhibition of mTORC1 by rapamycin was recently shown to attenuate the consolidation of extinction memories associated to an auditory threat. Moreover, an effector of mTOR, S6K1, was activated in the BLA during extinction, although apparently this activation was mediated by ERK (Huynh et al., 2017).

There was another correlation between gene expression in the amygdala and a behavioral variable in the extinction group, with *Gsk3α* expression being related to the number of lever presses during the first three days of extinction (Fig. 5). Self-administration extinction occurs during these three first days, such that this result could reflect a slower extinction rate in the animals that expressed more *Gsk3α* in the amygdala. GSK3β activity in the BLA is stronger after exposure to drug associated cues. Moreover, intraBLA injections of SB216763, a non-selective GSK3 inhibitor, immediately after the reactivation of cocaine cue memory impairs the preference of rats for the drug context cue, revealing a role for GSK3 in the persistence of the incentive value associated to drug associated cues (Wu et al., 2011). This effect was replicated in mice using i.p. SB216763 (Shi et al., 2014). Since SB216763 has a similar K_i_s for GSK3α and GSK3β (Coghlan et al., 2000), it is tempting to speculate that these effects could be mediated by either or both the GSK3 isoforms. Whether or not one of these isoforms in particular is responsible for the resistance to extinction of the response to drug associated cues, the administration of GSK3 inhibitors during withdrawal would seem to be an interesting approach for the treatment of addictions.

The only correlation involving a biochemical variable outside of the BLA was the inverse relationship between the amount of phPDK1 (Ser241) in the PFC and the slope of the regression line of morphine consumption during the first three days of extinction (Fig. 5). This behavioral variable can be considered an estimate of the extinction rate of the rats, indicating that the rats with more phPDK1 (Ser241) in the PFC extinguished faster. These results are coherent with the known activity of PDK1 and also, with the territories of the PFC dissected in our study. Indeed, PDK1 activity is required for Akt (Sarbassov et al., 2005), p70S6k (Pullen et al., 1998) and SGK1 activation (García-Martínez and Alessi, 2008), all part of the mTOR pathway and important regulators of protein synthesis. Alternatively, the PFC samples taken by us mostly contain the orbitofrontal cortex (OFC), a critical area signaling information about specific outcomes (Schoenbaum and Esber, 2010). Moreover, lesions or inactivation of the OFC impaired extinction of a reinforced response in rats (Joel and Klavir, 2006; Panayi and Killcross, 2014) and macaque monkeys (Butter, 1969; Izquierdo and Murray, 2005). In humans, extinction learning is also mediated by the OFC, as seen in a study using functional magnetic resonance imaging (Gottfried and Dolan, 2004).

## 5 Concluding remarks

In this study, we have addressed the putative effects of morphine self-administration and extinction on several elements of the mTOR pathway. Of the three areas studied, the most significant results were found in the amygdala. The role of this area in the processes of drug addiction and relapse is well known but to our knowledge, no one has previously observed the potential involvement of the mTOR pathway in this limbic structure. The genes and phosphoproteins identified are mainly involved in regulating protein synthesis, and they may also be recruited during memory formation and reconsolidation, concurring with earlier data. In the light of these findings, it would be interesting to more directly study the therapeutic value of this signaling pathway in opioid-related disorders.

## 6 Acknowledgements

This research was funded by the Spanish Ministerio de Economía y Competitividad (Project PSI2016-80541-P); the Ministerio de Sanidad, Servicios Sociales e Igualdad (Red de Trastornos Adictivos-Project RTA-RD16/020/0022 of the Instituto de Salud Carlos III; and the Plan Nacional sobre Drogas, Project 2016I073); the Dirección General de Investigación de la Comunidad de Madrid (Project S-2011/BMD-2308, Programa de Actividades I+D+I CANNAB-CM); the UNED (Plan de PromociÓn de la Investigación); and the European Union (Project JUST/2013/DPIP/AG/4823-EU MADNESS). We also thank Rosa Ferrado, Luis Carrillo, Gonzalo Moreno and Alberto Marcos for their excellent technical assistance.

## References

Bailey J, Ma D and Szumlinski KK (2012) Rapamycin attenuates the expression of cocaine-induced place preference and behavioral sensitization. Addiction Biology, Blackwell Publishing Ltd 17(2): 248–258. Available from: http://doi.wiley.com/10.1111/j.1369-1600.2010.00311.x (accessed 22 February 2017).

Beckley JT, Laguesse S, Phamluong K, et al. (2016) The First Alcohol Drink Triggers mTORC1-Dependent Synaptic Plasticity in Nucleus Accumbens Dopamine D1 Receptor Neurons. J. Neurosci. 36(3): 701–713.

Blommaart EFC, Luiken JJFP, Blommaart PJE, et al. (1995) Phosphorylation of ribosomal protein S6 is inhibitory for autophagy in isolated rat hepatocytes. Journal of Biological Chemistry 270(5): 2320–2326.

Brown AL, Flynn JR, Smith DW, et al. (2011) Down-regulated striatal gene expression for synaptic plasticity-associated proteins in addiction and relapse vulnerable animals. 2011/01/06. Int J Neuropsychopharmacol 14(8): 1099–1110. Available from: http://www.ncbi.nlm.nih.gov/entrez/query.fcgi?cmd=Retrieve&db=PubMed&dopt=Citation&list_uids=21205431.

Butter CM (1969) Perseveration in extinction and in discrimination reversal tasks following selective frontal ablations in Macaca mulatta. Physiology & Behavior 4(2): 163–171. Available from: http://www.sciencedirect.com/science/article/pii/0031938469900754 (accessed 15 June 2017).

Casadio A, Martin KC, Giustetto M, et al. (1999) A transient, neuron-wide form of CREB-mediated long-term facilitation can be stabilized at specific synapses by local protein synthesis. Cell 99(2): 221–37. Available from: http://www.ncbi.nlm.nih.gov/pubmed/10535740 (accessed 21 April 2017).

Chomczynski P and Sacchi N (1987) Single-step method of RNA isolation by acid guanidinium thiocyanate-phenol-chloroform extraction. Analytical Biochemistry 162(1): 156–159. Available from: http://linkinghub.elsevier.com/retrieve/pii/0003269787900212 (accessed 5 July 2017).

Coghlan MP, Culbert AA, Cross DA, et al. (2000) Selective small molecule inhibitors of glycogen synthase kinase-3 modulate glycogen metabolism and gene transcription. Chemistry & biology 7(10): 793–803. Available from: http://www.ncbi.nlm.nih.gov/pubmed/11033082 (accessed 10 June 2017).

Conover WJ (1999) Practical nonparametric statistics. Wiley.

Costa-Mattioli M, Sossin WS, Klann E, et al. (2009) Translational control of long-lasting synaptic plasticity and memory. Neuron, Cold Spring Harbor Laboratory Press, Cold Spring Harbor, NY 61(1): 10–26. Available from: http://www.ncbi.nlm.nih.gov/pubmed/19146809 (accessed 3 May 2017).

Cui Yue, Zhang XQ, Cui Y., et al. (2010) Activation of phosphatidylinositol 3-kinase/Akt-mammalian target of Rapamycin signaling pathway in the hippocampus is essential for the acquisition of morphine-induced place preference in rats. Neuroscience 171(1): 134–143. Available from: http://www.ncbi.nlm.nih.gov/pubmed/20826199 (accessed 18 January 2017).

Cunningham JT, Rodgers JT, Arlow DH, et al. (2007) mTOR controls mitochondrial oxidative function through a YY1-PGC-1alpha transcriptional complex. Nature 450(7170): 736–740.

Dayas C V, Smith DW and Dunkley PR (2012) An emerging role for the Mammalian target of rapamycin in ‘pathological’ protein translation: relevance to cocaine addiction. 2012/02/22. Front Pharmacol 3: 13. Available from: http://www.ncbi.nlm.nih.gov/entrez/query.fcgi?cmd=Retrieve&db=PubMed&dopt=Citation&list_uids=22347189.

Dos D. Sarbassov DD, Ali SM, Kim D-H, et al. (2004) Rictor, a Novel Binding Partner of mTOR, Defines a Rapamycin-Insensitive and Raptor-Independent Pathway that Regulates the Cytoskeleton. Current Biology 14(14): 1296–1302. Available from: http://www.ncbi.nlm.nih.gov/pubmed/15268862 (accessed 20 January 2017).

Düvel K, Yecies JL, Menon S, et al. (2010) Activation of a metabolic gene regulatory network downstream of mTOR complex 1. Molecular Cell 39(2): 171–183.

Everitt BJ, Cardinal RN, Parkinson JA, et al. (2003) Appetitive behavior: impact of amygdala-dependent mechanisms of emotional learning. 2003/05/02. Annals of the New York Academy of Sciences 985: 233–250. Available from: http://www.ncbi.nlm.nih.gov/entrez/query.fcgi?cmd=Retrieve&db=PubMed&dopt=Citation&list_uids=12724162.

Falls W, Miserendino M and Davis M (1992) Extinction of fear-potentiated startle: blockade by infusion of an NMDA antagonist into the amygdala. Journal of Neuroscience 12(3). Available from: http://www.jneurosci.org/content/12/3/854.long (accessed 6 June 2017).

Fuchs RA, Weber SM, Rice HJ, et al. (2002) Effects of excitotoxic lesions of the basolateral amygdala on cocaine-seeking behavior and cocaine conditioned place preference in rats. Brain Research 929(1): 15–25. Available from: http://www.sciencedirect.com/science/article/pii/S0006899301033662 (accessed 6 June 2017).

García-Martínez JM and Alessi DR (2008) mTOR complex 2 (mTORC2) controls hydrophobic motif phosphorylation and activation of serum-and glucocorticoid-induced protein kinase 1 (SGK1). Biochemical Journal 416(3): 375–385.

Gottfried JA and Dolan RJ (2004) Human orbitofrontal cortex mediates extinction learning while accessing conditioned representations of value. Nature Neuroscience 7(10): 1144–1152. Available from: http://www.nature.com/doifinder/10.1038/nn1314.

Guo N, Garcia MM and Harlan RE (2008) A morphine-paired environment alters c-Fos expression in the forebrain of rats displaying conditioned place preference or aversion. Behavioral Neuroscience, American Psychological Association 122(5): 1078–1086. Available from: http://doi.apa.org/getdoi.cfm?doi=10.1037/a0012595 (accessed 16 May 2017).

Hara K, Maruki Y, Long X, et al. (2002) Raptor, a binding partner of target of rapamycin (TOR), mediates TOR action. Cell 110(2): 177–89. Available from: http://www.ncbi.nlm.nih.gov/pubmed/12150926 (accessed 10 May 2017).

Hassan HE, Myers AL, Lee IJ, et al. (2009) Regulation of Gene Expression in Brain Tissues of Rats Repeatedly Treated by the Highly Abused Opioid Agonist, Oxycodone: Microarray Profiling and Gene Mapping Analysis. Drug Metabolism and Disposition 38(1). Available from: http://dmd.aspetjournals.org/content/38/1/157.long (accessed 24 May 2017).

Heller EA, Kaska S, Fallon B, et al. (2015) Morphine and cocaine increase serum-and glucocorticoid-inducible kinase 1 activity in the ventral tegmental area. Journal of Neurochemistry 132(2): 243–253.

Huynh TN, Santini E, Mojica E, et al. (2017) Activation of a novel p70 S6 kinase 1-dependent intracellular cascade in the basolateral nucleus of the amygdala is required for the acquisition of extinction memory. Molecular Psychiatry. Available from: http://www.ncbi.nlm.nih.gov/pubmed/28461701 (accessed 9 June 2017).

Izquierdo A and Murray EA (2005) Opposing effects of amygdala and orbital prefrontal cortex lesions on the extinction of instrumental responding in macaque monkeys. European Journal of Neuroscience 22(9): 2341–2346. Available from: http://www.ncbi.nlm.nih.gov/pubmed/16262672 (accessed 15 June 2017).

James MH, Quinn RK, Ong LK, et al. (2016) Rapamycin reduces motivated responding for cocaine and alters GluA1 expression in the ventral but not dorsal striatum. European Journal of Pharmacology 784: 147–154.

Jefferies HBJ, Reinhard C, Kozma SC, et al. (1994) Rapamycin selectively represses translation of the ‘polypyrimidine tract’ mRNA family. Cell Biology 91: 4441–4445. Available from: http://www.pnas.org/content/91/10/4441.full.pdf (accessed 11 June 2017).

Joel D and Klavir O (2006) The effects of temporary inactivation of the orbital cortex in the signal attenuation rat model of obsessive compulsive disorder. Behavioral Neuroscience 120(4): 976–983. Available from: http://www.ncbi.nlm.nih.gov/pubmed/16893303 (accessed 15 June 2017).

Kalivas PW and O’Brien C (2008) Drug addiction as a pathology of staged neuroplasticity. Neuropsychopharmacology 33(1): 166–180. Available from: http://www.ncbi.nlm.nih.gov/entrez/query.fcgi?cmd=Retrieve&db=PubMed&dopt=Citation&list_uids=17805308.

Kauer JA and Malenka RC (2007) Synaptic plasticity and addiction. Nature reviews. Neuroscience, Nature Publishing Group 8(11): 844–58. Available from: http://dx.doi.org/10.1038/nrn2234 (accessed 12 March 2015).

Kim DH and Sabatini DM (2004) Raptor and mTOR: subunits of a nutrient-sensitive complex. Current topics in microbiology and immunology 279: 259–70. Available from: http://www.ncbi.nlm.nih.gov/pubmed/14560962 (accessed 10 May 2017).

Kimball SR, Shantz LM, Horetsky RL, et al. (1999) Leucine regulates translation of specific mRNAs in L6 myoblasts through mTOR-mediated changes in availability of eIF4E and phosphorylation of ribosomal protein S6. The Journal of biological chemistry, American Society for Biochemistry and Molecular Biology 274(17): 11647–52. Available from: http://www.ncbi.nlm.nih.gov/pubmed/10206976 (accessed 11 June 2017).

Kwon C-H, Zhu X, Zhang J, et al. (2003) mTor is required for hypertrophy of Pten-deficient neuronal soma in vivo. Proceedings of the National Academy of Sciences 100(22): 12923–12928. Available from: http://www.ncbi.nlm.nih.gov/pubmed/14534328 (accessed 20 January 2017).

Ladner CL, Yang J, Turner RJ, et al. (2004) Visible fluorescent detection of proteins in polyacrylamide gels without staining. Analytical Biochemistry 326(1): 13–20. Available from: http://linkinghub.elsevier.com/retrieve/pii/S0003269703008017 (accessed 5 July 2017).

Law PY, Wong YH and Loh HH (2000) Molecular mechanisms and regulation of opioid receptor signaling. Annual review of pharmacology and toxicology 40(13): 389–430.

Lee CT, Tyan SW, Ma YL, et al. (2006) Serum-and glucocorticoid-inducible kinase (SGK) is a target of the MAPK/ERK signaling pathway that mediates memory formation in rats. European Journal of Neuroscience 23(5): 1311–1320.

Li Y, Eitan S, Wu J, et al. (2003) Morphine Induces Desensitization of Insulin Receptor Signaling. Molecular and Cellular Biology 23(17): 6255–6266.

Lin J, Liu L, Wen Q, et al. (2014) Rapamycin prevents drug seeking via disrupting reconsolidation of reward memory in rats. International Journal of Neuropsychopharmacology 17(1): 127–136. Available from: http://ovidsp.ovid.com/ovidweb.cgi?T=JS&CSC=Y&NEWS=N&PAGE=fulltext&D=psyc11&AN=2013-43199-012%5Cnhttp://mcgill.on.worldcat.org/atoztitles/link?sid=OVID:psycdb&id=pmid:&id=10.1017%2FS1461145713001156&issn=1461-1457&isbn=&volume=17&issue=1&spage=127&pag.

Liu-Yesucevitz L, Bassell GJ, Gitler AD, et al. (2011) Local RNA Translation at the Synapse and in Disease. Journal of Neuroscience 31(45): 16086–16093. Available from: http://www.ncbi.nlm.nih.gov/pubmed/22072660 (accessed 3 May 2017).

Livak KJ and Schmittgen TD (2001) Analysis of Relative Gene Expression Data Using Real-Time Quantitative PCR and the 2???CT Method. Methods 25(4): 402–408. Available from: http://linkinghub.elsevier.com/retrieve/pii/S1046202301912629 (accessed 16 June 2017).

Lu K-T, Walker DL and Davis M (2001) Mitogen-Activated Protein Kinase Cascade in the Basolateral Nucleus of Amygdala Is Involved in Extinction of Fear-Potentiated Startle. Journal of Neuroscience 21(16). Available from: http://www.jneurosci.org/content/21/16/RC162.long (accessed 6 June 2017).

Luo Y-X, Xue Y-X, Shen H-W, et al. (2013) Role of amygdala in drug memory. Neurobiology of Learning and Memory 105: 159–173. Available from: http://www.sciencedirect.com/science/article/pii/S1074742713001093 (accessed 16 May 2017).

Lüscher C and Malenka RC (2011) Drug-Evoked Synaptic Plasticity in Addiction: From Molecular Changes to Circuit Remodeling. Neuron.

Ma XM and Blenis J (2004) Molecular mechanisms of mTOR-mediated translational control. Nature Reviews: Molecular Cell Biology 5(10): 827–835.

Ma XM and Blenis J (2009) Molecular mechanisms of mTOR-mediated translational control. Nature Reviews Molecular Cell Biology, Nature Publishing Group 10(5): 307–318. Available from: http://www.nature.com/doifinder/10.1038/nrm2672 (accessed 12 May 2017).

Ma YL (2006) SGK protein kinase facilitates the expression of long-term potentiation in hippocampal neurons. Learning & Memory 13(2): 114–118.

Mateyak MK and Kinzy TG (2010) eEF1A: Thinking outside the ribosome. Journal of Biological Chemistry.

Mazei-Robison MS, Koo JW, Friedman AK, et al. (2011) Role for mTOR signaling and neuronal activity in morphine-induced adaptations in ventral tegmental area dopamine neurons. 2011/12/27. Neuron 72(6): 977–990. Available from: http://www.ncbi.nlm.nih.gov/entrez/query.fcgi?cmd=Retrieve&db=PubMed&dopt=Citation&list_uids=22196333.

McLellan AT, Lewis DC, O’Brien CP, et al. (2000) Drug Dependence, a Chronic Medical Illness. JAMA, American Medical Association 284(13): 1689. Available from: http://jama.jamanetwork.com/article.aspx?doi=10.1001/jama.284.13.1689 (accessed 20 April 2017).

Narita M, Akai H, Kita T, et al. (2005) Involvement of mitogen-stimulated p70-S6 kinase in the development of sensitization to the methamphetamine-induced rewarding effect in rats. Neuroscience 132(3): 553–560.

Neasta J, Barak S, Hamida S Ben, et al. (2014) mTOR complex 1: a key player in neuroadaptations induced by drugs of abuse. Journal of neurochemistry 130(2): 172–84.

Niswender KD, Gallis B, Blevins JE, et al. (2003) Immunocytochemical Detection of Phosphatidylinositol 3-kinase Activation by Insulin and Leptin. Journal of Histochemistry & Cytochemistry 51(3): 275–283. Available from: http://www.ncbi.nlm.nih.gov/pubmed/12588955 (accessed 8 May 2017).

Panayi MC and Killcross S (2014) Orbitofrontal cortex inactivation impairs between-but not within-session Pavlovian extinction: An associative analysis. Neurobiology of Learning and Memory 108: 78–87. Available from: http://www.ncbi.nlm.nih.gov/pubmed/23954805 (accessed 16 June 2017).

Pearce LR, Komander D and Alessi DR (2010) The nuts and bolts of AGC protein kinases. Nature reviews. Molecular cell biology 11(1): 9–22. Available from: http://www.nature.com/doifinder/10.1038/nrm2822 (accessed 20 January 2017).

Polakiewicz RD, Schieferl SM, Gingras AC, et al. (1998) mu-Opioid receptor activates signaling pathways implicated in cell survival and translational control. The Journal of biological chemistry 273(36): 23534–41. Available from: http://www.ncbi.nlm.nih.gov/pubmed/9722592 (accessed 17 January 2017).

Porstmann T, Santos CR, Griffiths B, et al. (2008) SREBP Activity Is Regulated by mTORC1 and Contributes to Akt-Dependent Cell Growth. Cell Metabolism 8(3): 224–236.

Pullen N, Dennis PB, Andjelkovic M, et al. (1998) Phosphorylation and activation of p70s6k by PDK1. Science (New York, N.Y.) 279(5351): 707–10. Available from: http://www.ncbi.nlm.nih.gov/pubmed/9445476 (accessed 15 June 2017).

Richter JD and Sonenberg N (2005) Regulation of cap-dependent translation by eIF4E inhibitory proteins. Nature 433(7025): 477–480. Available from: http://www.ncbi.nlm.nih.gov/pubmed/15690031 (accessed 17 May 2017).

Ruijter JM, Ramakers C, Hoogaars WMH, et al. (2009) Amplification efficiency: linking baseline and bias in the analysis of quantitative PCR data. Nucleic Acids Research 37(6): e45–e45. Available from: http://www.ncbi.nlm.nih.gov/pubmed/19237396 (accessed 23 January 2017).

Russo SJ, Bolanos CA, Theobald DE, et al. (2007) IRS2-Akt pathway in midbrain dopamine neurons regulates behavioral and cellular responses to opiates. 2006/12/05. Nature Neuroscience 10(1): 93–99. Available from: http://www.ncbi.nlm.nih.gov/entrez/query.fcgi?cmd=Retrieve&db=PubMed&dopt=Citation&list_uids=17143271.

Sarbassov DD, Guertin DA, Ali SM, et al. (2005) Phosphorylation and Regulation of Akt/PKB by the Rictor-mTOR Complex. Science 307(5712). Available from: http://science.sciencemag.org/content/307/5712/1098.full (accessed 15 June 2017).

Schieke SM, Phillips D, McCoy JP, et al. (2006) The mammalian target of rapamycin (mTOR) pathway regulates mitochondrial oxygen consumption and oxidative capacity. The Journal of biological chemistry 281(37): 27643–52. Available from: http://www.ncbi.nlm.nih.gov/pubmed/16847060.

Schoenbaum G and Esber GR (2010) How do you (estimate you will) like them apples? Integration as a defining trait of orbitofrontal function. Current opinion in neurobiology, NIH Public Access 20(2): 205–11. Available from: http://www.ncbi.nlm.nih.gov/pubmed/20206497 (accessed 15 June 2017).

Schroeder JP and Packard MG (2004) Facilitation of Memory for Extinction of Drug-Induced Conditioned Reward: Role of Amygdala and Acetylcholine. Learning & Memory, Cold Spring Harbor Laboratory Press 11(5): 641–647. Available from: http://www.ncbi.nlm.nih.gov/pubmed/15466320 (accessed 6 June 2017).

Shi J, Jun W, Zhao L-Y, et al. (2009) Effect of rapamycin on cue-induced drug craving in abstinent heroin addicts. European Journal of Pharmacology 615(1–3): 108–112. Available from: http://www.ncbi.nlm.nih.gov/pubmed/19470385 (accessed 18 January 2017).

Shi X, Miller JS, Harper LJ, et al. (2014) Reactivation of cocaine reward memory engages the Akt/GSK3/mTOR signaling pathway and can be disrupted by GSK3 inhibition. Psychopharmacology, Springer Berlin Heidelberg 231(16): 3109–3118. Available from: http://link.springer.com/10.1007/s00213-014-3491-8 (accessed 10 June 2017).

Shimobayashi M and Hall MN (2014) Making new contacts: the mTOR network in metabolism and signalling crosstalk. Nature reviews. Molecular cell biology, Nature Publishing Group 15(3): 155–62. Available from: http://www.ncbi.nlm.nih.gov/pubmed/24556838.

Singh P, Sharma B, Gupta S, et al. (2015) In vivo and in vitro attenuation of naloxone-precipitated experimental opioid withdrawal syndrome by insulin and selective K<inf>ATP</inf> channel modulator. Psychopharmacology 232(2): 465–475.

Stoica L, Zhu PJ, Huang W, et al. (2011) Selective pharmacogenetic inhibition of mammalian target of Rapamycin complex I (mTORC1) blocks long-term synaptic plasticity and memory storage. Proceedings of the National Academy of Sciences of the United States of America 108(9): 3791–3796. Available from: http://www.ncbi.nlm.nih.gov/pubmed/21307309%5Cnhttp://www.pnas.org/content/108/9/3791.full.pdf+html. Stripping for reprobing (n.d.).

Taha C and Klip A (1999) The Insulin Signaling Pathway. The Journal of Membrane Biology, Springer-Verlag 169(1): 1–12. Available from: http://link.springer.com/10.1007/PL00005896 (accessed 8 May 2017).

Tsai KJ, Chen SK, Ma YL, et al. (2002) *sgk*, a primary glucocorticoid-induced gene, facilitates memory consolidation of spatial learning in rats. Proceedings of the National Academy of Sciences 99(6): 3990–3995.

Tsokas P, Grace EA, Chan P, et al. (2005) Local Protein Synthesis Mediates a Rapid Increase in Dendritic Elongation Factor 1A after Induction of Late Long-Term Potentiation. Journal of Neuroscience 25(24). Available from: http://www.jneurosci.org/content/25/24/5833 (accessed 11 June 2017).

Wang X, Luo Y -x., He Y -y., et al. (2010) Nucleus Accumbens Core Mammalian Target of Rapamycin Signaling Pathway Is Critical for Cue-Induced Reinstatement of Cocaine Seeking in Rats. Journal of Neuroscience 30(38): 12632–12641. Available from: http://www.ncbi.nlm.nih.gov/pubmed/20861369 (accessed 23 February 2017).

Wu J, McCallum SE, Glick SD, et al. (2011) Inhibition of the mammalian target of rapamycin pathway by rapamycin blocks cocaine-induced locomotor sensitization. Neuroscience 172: 104–109. Available from: http://www.ncbi.nlm.nih.gov/pubmed/20977929 (accessed 22 February 2017).

Wu P, Xue Y, Ding Z, et al. (2011) Glycogen synthase kinase 3? in the basolateral amygdala is critical for the reconsolidation of cocaine reward memory. Journal of Neurochemistry, Blackwell Publishing Ltd 118(1): 113–125. Available from: http://doi.wiley.com/10.1111/j.1471-4159.2011.07277.x (accessed 10 June 2017).

Zhou J, Blundell J, Ogawa S, et al. (2009) Pharmacological inhibition of mTORC1 suppresses anatomical, cellular, and behavioral abnormalities in neural-specific Pten knock-out mice. The Journal of neuroscience: the official journal of the Society for Neuroscience 29(6): 1773–83. Available from: http://www.jneurosci.org/cgi/doi/10.1523/JNEUROSCI.5685-08.2009 (accessed 20 January 2017).

